# Paired SARS-CoV-2 Spike Protein Mutations Observed During Ongoing SARS-CoV-2 Viral Transfer from Humans to Minks and Back to Humans

**DOI:** 10.1101/2020.12.22.424003

**Authors:** Scott Burkholz, Suman Pokhrel, Benjamin R. Kraemer, Daria Mochly-Rosen, Richard T. Carback, Tom Hodge, Paul Harris, Serban Ciotlos, Lu Wang, CV Herst, Reid Rubsamen

**Author notes:** Corresponding Author, Dr. Reid Rubsamen, M.D., Department of Anesthesiology and Perioperative Medicine, University Hospitals, Cleveland, Medical Center, Case Western Reserve School of Medicine, 11100 Euclid Ave, Cleveland, OH, 44106.

## Abstract

A mutation analysis of SARS-CoV-2 genomes collected around the world sorted by sequence, date, geographic location, and species has revealed a large number of variants from the initial reference sequence in Wuhan. This analysis also reveals that humans infected with SARS-CoV-2 have infected mink populations in the Netherlands, Denmark, United States, and Canada. In these animals, a small set of mutations in the spike protein receptor binding domain (RBD), often occurring in specific combinations, has transferred back into humans. The viral genomic mutations in minks observed in the Netherlands and Denmark show the potential for new mutations on the SARS-CoV-2 spike protein RBD to be introduced into humans by zoonotic transfer. Our data suggests that close attention to viral transfer from humans to farm animals and pets will be required to prevent build-up of a viral reservoir for potential future zoonotic transfer.

## 1. Introduction

Coronaviruses are thought to have ancient origins extending back tens of millions of years with coevolution tied to bats and birds ^1^. This subfamily of viruses contains proofreading mechanisms, rare in other RNA viruses, reducing the frequency of mutations that might alter viral fitness ^2^. The D614G mutation, which lies outside the RBD, is an example of a fitness-enhancing mutation on the spike glycoprotein that became the most prevalent variant as the virus spreads through human populations ^3^. The recently observed, and more infectious, D796H mutation paired with ΔH69/V70, also outside the RBD domain, first observed in January 2020, has spread throughout Southeast England ^4^. Data from next-generation sequencing has shown that the SARS-CoV-2 viral genome mutates at about half the rate of Influenza and about a quarter of the rate seen for HIV, with about 10 nucleotides of average difference between samples^5^. The Global Initiative on Sharing Avian Influenza Data (GISAID)^6,7^, has catalogued over 235,299 SARS-CoV-2 sequences to date from samples provided by laboratories around the world. This diversity is profound with well over 12,000 mutations having been shown to exist, with the potential for non-synonymous substitutions, insertions, or deletions resulting in amino acid changes which could result in structural and functional changes in virus proteins ^5,8–10^. While Coronaviruses initially developed in animals and transferred to humans, transfer back to animals and then back to humans again has recently been observed in in the *Neovison vison* species of mink, currently being raised in farms around the world.

Non-silent mutations in the SARS-CoV-2 spike glycoprotein gene can produce structural and functional changes impacting host receptor binding, and viral entry into the cell. Specifically, a mutation in the RBD region, residues 333-527 ^11^ can potentially either increase or decrease the affinity of spike protein for the human ACE2 receptor, directly influencing the ability of the virus to enter a cell and successfully infect the host. The receptor binding motif (RBM), located within the RBD from 438 to 506 ^11^ plays a key role in enabling virus contact with ACE2 and is an optimal target for neutralizing antibodies ^12^. Mutations in this region, which also corresponds to the complementarity-determining regions (CDR) of a potentially binding immunoglobulin, could affect the antibody’s ability to neutralize the virus ^13^.

Here we examined the mutations that were associated with the transfer of SARS-CoV-2 from humans to minks and back to humans. This example of zoonotic transfer highlights the importance of paired mutations in the RBD domain and suggests potential challenges for sustained efficacy of neutralizing antibodies focusing on that region.

## 2. Methods

### 2.1 Viral Sequencing Analysis

All 235,299 available spike glycoprotein amino acid sequences available thru December 5^th^ were downloaded from GISAID^6,7^. Before downloading, full genome nucleotide sequences were individually aligned via MAFFT^14^ to the WIV04 (MN996528.1)^15^ internal reference sequence. Aligned sequences were translated to amino acids based on spike glycoprotein reference sequence positions at 21563 to 25384.

The downloaded spike glycoprotein file was re-aligned via MAFFT ^14^ against WIV04 (MN996528.1)^15^ with position numbering kept constant. Sequences were split between human (206,591 sequences) and the *Neovison vison* species of mink (332 sequences).

A custom Python ^16^ script was utilized to compare the variants seen in mink and humans differing from the WIV04 reference sequence. Only mutations with at least three occurrences in minks were retained. After initial examination of the whole genome, all mutations inside of, or within 25 amino acids of, RBD (333-527)^11^ were also kept for analysis. The script was rerun using only sequences and meta-data matching those criteria. Thirteen human and three mink sequences were removed that were found to contain 25 or more gaps or mutations compared to the WIV04 reference. This was done to avoid the potential for improper mutation identification confounding downstream analysis. After all these filtering procedures, 782 human and 251 mink sequences remained for analysis.

The statistical package R ^17^ was utilized to plot mutation frequency by geography, date, and species to reveal patterns indicative of zoonotic transfer. Sequence identifiers, and the respective authors, utilized for results are shown in supplementary table 1.

### 2.2 SARS Alignment Comparisons

The SARS-CoV-2 reference sequence WIV04 (MN996528.1)^15^ was aligned against a SARS-CoV-1 reference sequence (NC_004718.3)^18^ via MAFFT^14^. Residue positions were visualized in Jalview 19

### 2.3 3D Structure Visualization

The PDB file, 7A98 ^20^, was downloaded from RCSB.org. Positions of interest were visualized in MOE ^21^ for figure 1b.

**Figure 1.**
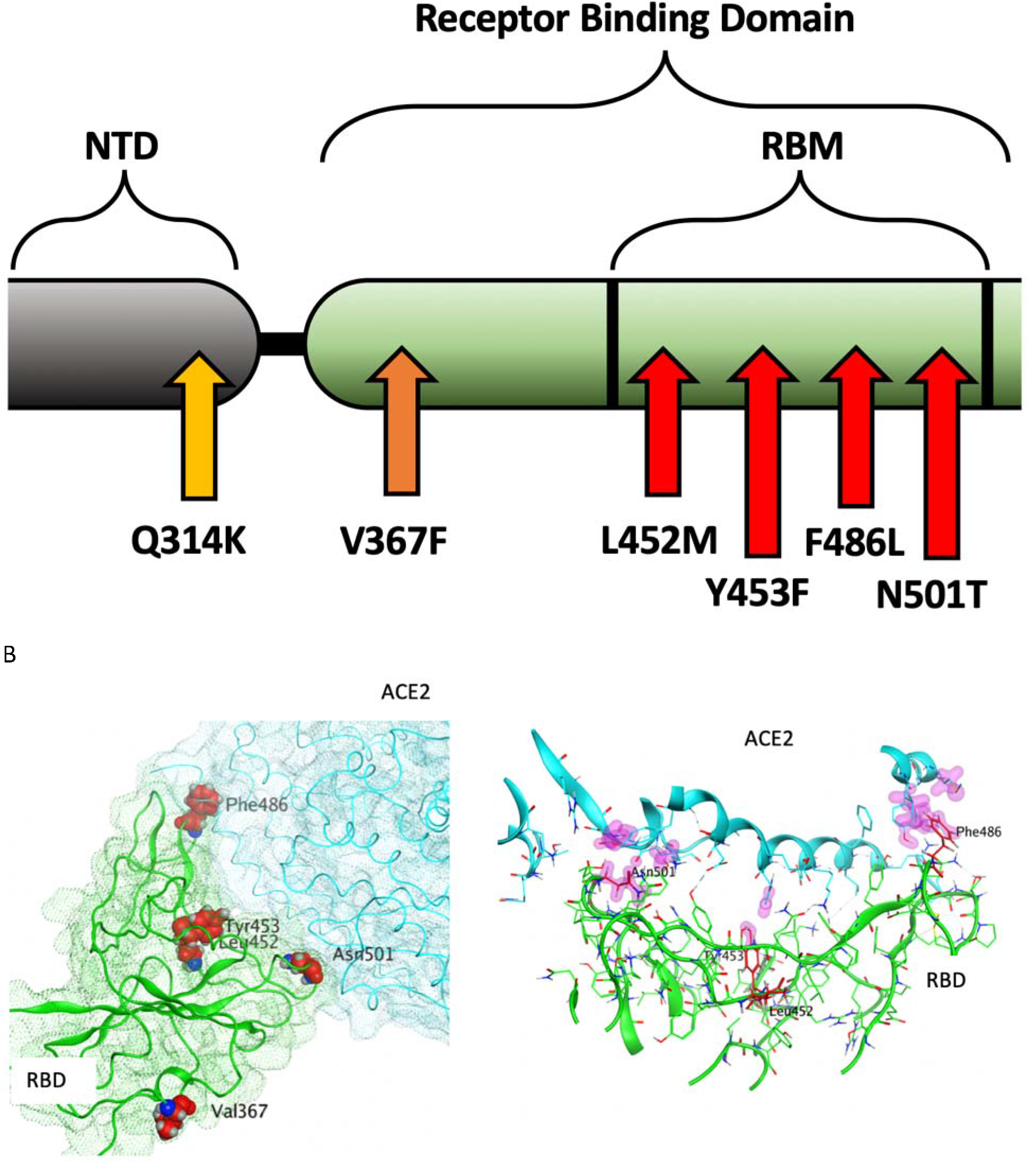
A: Illustration of positions of mutation variants described on the SARS-CoV-2 spike glycoprotein for the N-terminal domain (NTD), receptor binding domain (RBD), and receptor binding motif (RBM). B: Left: Crystal Structure (PDB ID: 7A98) of SARS-CoV-2 receptor binding domain (green) in complex with ACE2 (teal) with residues highlighted in red. Right: Interaction of highlighted residues (red) with ACE2 (teal). Interaction denoted with magenta clouds.

### 2.4 Molecular Effect Calculations

Molecular Operating Environment (MOE)^21^ software was used with PDB 7A98 ^20^, and prepared with QuickPrep functionality at the default settings, to optimize the H-bond network and perform energy minimization on the system. Affinity calculations were performed using 7A98.A (spike protein monomer) and 7A98.D (ACE2) chains. Residues in spike protein (7A98.A) within 4.5 Å from ACE2 (7A98.D) were selected and the residue scan application was run by defining ACE2 (7A98.D) as the ligand. Residue scans and change in affinity calculations were also performed on L452. Stability calculations were performed by running the residue scan application using residues in RBD (331-524) of spike protein (7A98.A). Residue scans and change in stability calculations were also performed on Q314. The changes in stability and affinity (kcal/mol) between the variants and the reference sequence were calculated as per MOE’s definition ^21^. The potential effect of variants was predicted using PROVEAN ^22^ with R^17^, and SIFT ^23^. Residues in ACE2 protein (7A98.D) within 4.5 Å from spike protein (7A98.A) were selected and the residue scan application was run by defining the spike protein (7A98.A) as the ligand. The changes in affinity (kcal/mol) between mink, mouse, and hamster ACE2 sequences compared to human ACE2 were calculated as per MOE’s definition^21^.

### 2.5 Phylogenic Tree

IQ-TREE 2 ^24^ was used to generate a phylogenic tree via maximum likelihood calculations. The RBD region, plus 25 amino acids on each end, of the filtered, processed human and mink samples was imputed for analysis. The “FLU+I” model was chosen as the best model to fit, with subsequent ultrafast bootstrapping till convergence and branch testing. FigTree (http://tree.bio.ed.ac.uk/software/figtree/) was used to visualize and color the phylogenic tree.

### 2.6 ACE2 sequence alignment

ACE2 protein sequences were obtained from Uniprot^25^ for human (identifier: Q9BYF1-1), mouse (identifier: Q8R0I0-1), hamster (identifier: (A0A1U7QTA1-1), and from the NCBI protein database for mink (identifier: QPL12211.1). The alignment was visualized in Jalview ^19^ and colored according to sequence identity. The similarity scores for the entire protein sequence and the spike receptor binding domain motif were calculated in MOE^21^. Residues in ACE2 within 4.5Å of spike receptor binding domain were defined as the receptor binding motif.

## 3. Results

The following six mutations met all the criteria described in the methods section and demonstrate the potential for zoonotic transfer between species. Mutations outside of the spike RBD and other proteins were examined but did not result in significant findings.

### 3.1 Netherlands

#### F486L

The F486L mutation has a substitution of leucine for phenylalanine occurring within the RBM on the SARS-CoV-2 spike glycoprotein (figure 1 A, B). These amino acids are similar in physiochemical properties, with an aromatic ring being replaced by an aliphatic chain. This new variant in F486L, conserved across SARS-CoV-1 and SARS-CoV-2, was first seen via the strain RaTG13 in a *Rhinolophus affinis* bat sample collected in Yunnan, China during 2013. In 2017, this variant was also found in *Manis javanica*, a species of pangolin. F486L was not present in human sequences from the dataset at the start of the pandemic, but started to appear in minks at the end of April 2020. We found 125 sequences from mink samples with this mutation collected in the Netherlands since that time.

A sample submission date places the first known potential transfer back from minks to humans in August 2020, also in the Netherlands. Although sample collection dates are unavailable for these human samples, submission dates show a larger number of human samples with this variant were reported in the Netherlands in October and November 2020. One human sample in Scotland, collected in October 2020 shows that F486L may be viable alone and can occur *de novo*, without evidence of zoonotic transfer. Based on mutations seen with F486L, L452M and Q314K, and considering the potential for sequencing error, this case is likely not linked to the Netherlands sequences (table 1).

**Table 1:** Mutations counted by species, location, and linkage with other mutations. Highlighted mutations illustrate inheritance patterns.

| Country | Mutations | Human Count | Mink Count |
| --- | --- | --- | --- |
| F486L |  |  |  |
| Netherlands | F486L, L452M | 6 | 32 |
| Netherlands | F486L, Q314K | 13 | 84 |
| Other (Michigan) | F486L, N501T | 2 | N/A |
| Other | F486L | 1 | N/A |
| Netherlands | F486L | 0 | 9 |
| L452M |  |  |  |
| Netherlands | L452M, F486L | 6 | 32 |
| Denmark | L452M | 1 | 0 |
| Other | L452M | 4 | N/A |
| Q314K |  |  |  |
| Netherlands | Q314K, F486L | 13 | 84 |
| Other | Q314K | 10 | N/A |
| Y453F |  |  |  |
| Denmark | Y453F | 629 | 85 |
| Netherlands | Y453F | 4 | 32 |
| Netherlands | Y453F, V367F | 1 | 1 |
| Other | Y453F | 5 | N/A |
| N501T |  |  |  |
| Other (Michigan) | N501T, F486L | 2 | N/A |
| Other | N501T | 10 | N/A |
| Netherlands | N501T | 0 | 4 |
| V367F |  |  |  |
| Netherlands | V367F, Y453F | 1 | 1 |
| Netherlands | V367F | 9 | 4 |
| Denmark | V367F | 2 | 0 |
| Other | V367F | 85 | N/A |

The L452M and Q314K variants were almost always observed to appear concurrently with F486L, apart from the first few known mink sequences (figure 2A). The only other sequences in the database with these variants outside the Netherlands were from two human samples collected in Michigan, coinciding with an October 2020 mink farm outbreak there ^26^. These latter human samples with F486L contain a second variant, N501T, which were also observed in four mink samples obtained in the Netherlands. The findings of some mutations with others (figure 3) suggests the potential requirement of a specific second mutation, L452M or Q314K, to be present in order for the virus to jump back from minks to humans.

**Figure 2:**
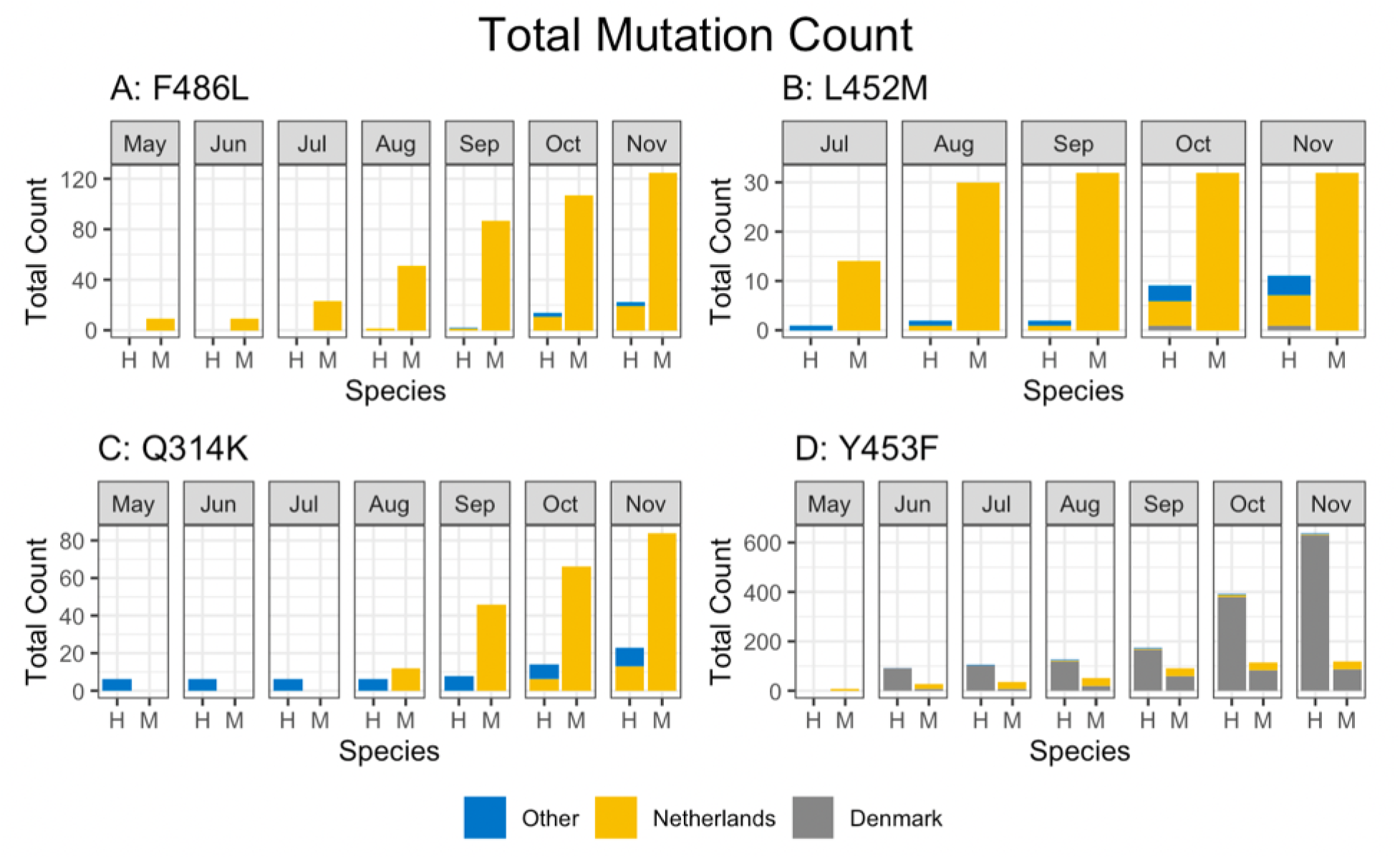
Total mutation counts stratified by mutation, location, and species (human and mink). Dates reported for human samples from the Netherlands are predominately submission dates, in contrast to the reported collection dates seen in meta data associated with sequences submitted from other countries.

**Figure 3:**
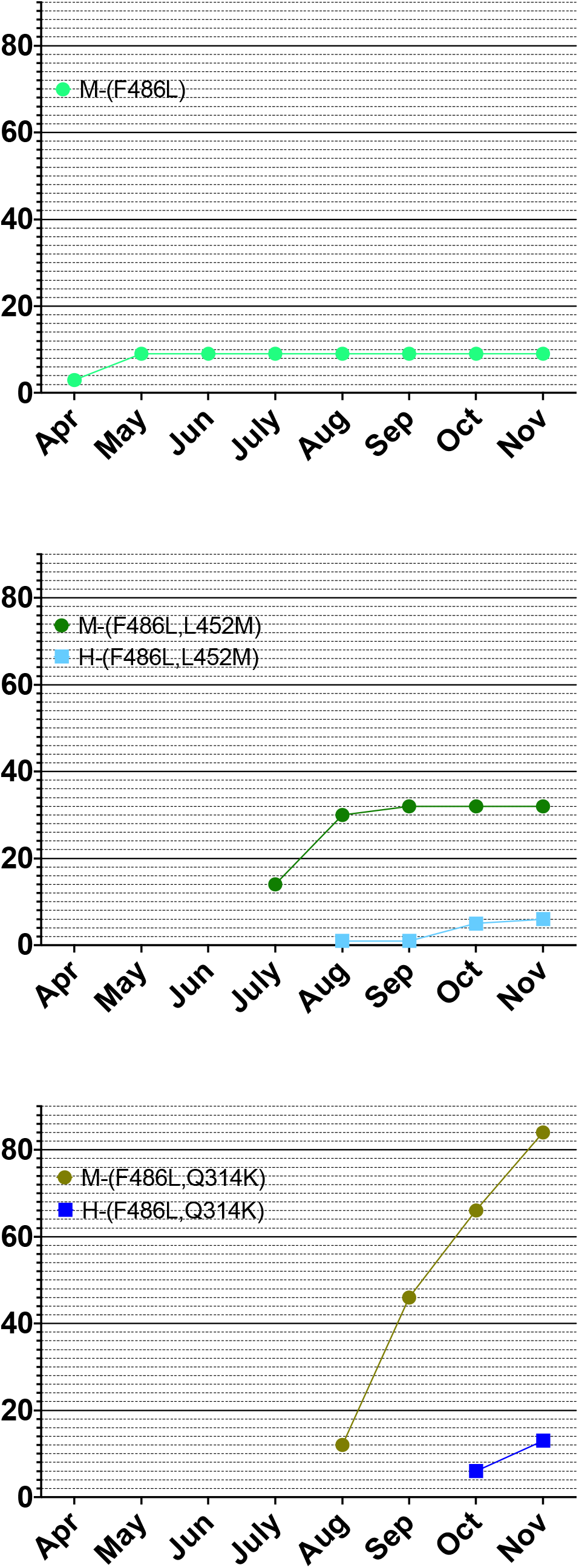
Cumulative counts for mutation F486L individually and in pairs with L452M or Q314K in the Netherlands.

#### L452M

The L452M variant in the spike RBM involves substitution of methionine for leucine at position 452 (figure 1 A, B). The side chains for these amino acids are quite similar, with the addition of a sulfur. This variant appeared first in mink samples from the dataset dated July 2020 and was found to co-occur with F486L. We also observed that L452M and F486L occurred together in all human samples to date from the Netherlands (table 1), submitted between August 2020 and through November 2020. The five human samples containing L452M that were collected outside the Netherlands were not paired with F486L in the sequence database (figure 2B), suggesting independent emergence.

#### Q314K

Q314K is located close to the RBD, and has lysine substituted for glutamine at position 314 (figure 1A) resulting in the addition of a positive charge to the side chain. This new variant in Q314K is found in both SARS-CoV-1 and SARS-CoV-2. The earliest sequences in the database with this mutation were found in five human samples taken in Northern California and one human sample taken in Mexico in May 2020. This variant first appeared in human samples submitted from the Netherlands in October and November 2020 (figure 2C). The dataset showed that Q314K was first seen in minks starting in August 2020 and was always observed to occur with F486L, but only in the absence of L452M. The ten human samples from outside the Netherlands all had Q314K without the other variants described here (table 1), suggesting it can occur *de-novo*.

### 3.2 Denmark

#### Y453F

Y453F is an RBM mutation substituting phenylalanine for tyrosine at position 453 that we observed present in a substantial number of human sequences in the database from samples submitted from around the world since the start of the pandemic (figure 1A, B). The substituted amino acid is similar to the reference with the change being less polar. This new variant in Y453F is found in both SARS-CoV-1 and SARS-CoV-2. The sequences analyzed for this study show this mutation appearing in minks starting in the Netherlands in April, and Denmark in June 2020. We saw rapid expansion of this mutation, present in 629 human sample sequences from Denmark, beginning in June 2020 (figure 2D, table 1). Of the five samples collected outside the Netherlands or Denmark, one sample came from Utah, one of the top producers of mink fur in the United States and that was experiencing a SARS-CoV-2 outbreak on mink farms at the same time^27^.

### 3.3 United States

#### N501T

N501T is a mutation in the RBM and involves a change from asparagine to threonine at position 501 (figure 1A, B). This new N501T variant is found in both SARS-CoV-1 and SARS-CoV-2 and was first described in 2017 in a species of pangolin, where it was seen to co-occur with F486L. N501T was found only sixteen times in humans or minks in the samples we analyzed. Four mink samples with N501T were submitted from the Netherlands between April and June 2020. The twelve human sequences with the N501T variant were seen in samples collected from March to October 2020 but were not seen in samples collected in the Netherlands or Denmark (table 1). As mentioned previously, two of the three sequences among the twelve collected outside the Netherlands with F486L contain this variant and were from Michigan, a state with documented mink farm outbreaks. Two of the twelve human sequences with N501T alone were found in Wisconsin on October 3^rd^. This variant was present in a human sample collected in Taylor county, where a mink farm was reported on October 8^th^ to have confirmed cases, now with a second farm outbreak and over 5,400 mink deaths from the virus ^28,29^.

#### V367F

V367F at the RBD, position 367 (figure 1 A, B), results in addition an aromatic group (substituting phenylalanine for valine). This new variant in V367F is found in both SARS-CoV-1 and SARS-CoV-2. Our analysis identified this variant in 97 human samples collected around the world, not localized to any one region (table 1). V367F was present in five minks and ten humans in samples collected in the Netherlands. We observed that this variant can occur *de-novo* without zoonotic transfer. Some cases in humans were seen in samples collected between August and October 2020, in Oregon, another state that was experiencing a mink farm outbreak at the same time^30^. We found nine human samples with this variant in Washington State between February and August; although there were no reports on SARS-CoV-2 outbreak in minks here, there are a small number of farms ^31^.

### 3.4 Mutation Effects

The six mutations we studied result from a Single Nucleotide Polymorphism (SNP), and are among those with the least potential consequence on the stability (kcal/mol, supplementary table 2) of the spike glycoprotein. Analysis for human ACE2 binding affinity (kcal/mol, supplementary table 3) determined that F486L, Y453F, N501T, and L452M likely cause minimal effects on the affinity of the spike protein to bind ACE2. The calculated difference in affinity score for the Y453F mutation indicates a slightly increased binding potential of this spike variant for hACE2 as compared to the reference amino acid sequence. Predicted effects of all six mutations illustrate neutral and tolerated changes (supplementary table 4) suggesting that the function of the protein is not drastically changed, and likely remains fit for replication and infection.

### 3.5 Phylogenic Tree

To examine the statistical relationships between variant sequences, a phylogeny via maximum likelihood estimation was created. Sequences with one of the described mutations were utilized by taking the protein sequences of the RBD, with an additional 25 amino acids on either side. Visualization reveals distinct grouping by mutation category (figure 4). These groupings further show that these mutations can occur individually or together. Human and mink sequences also group together, illustrating the similarities for SARS-CoV-2 between these two species. Once these mutations are present, additional mutations do not appear to be required to facilitate virus movement between humans and minks. A plot without these distinct groupings from the identified mutation categories in both humans and minks, could indicate additional variants positions present in samples.

**Figure 4:**
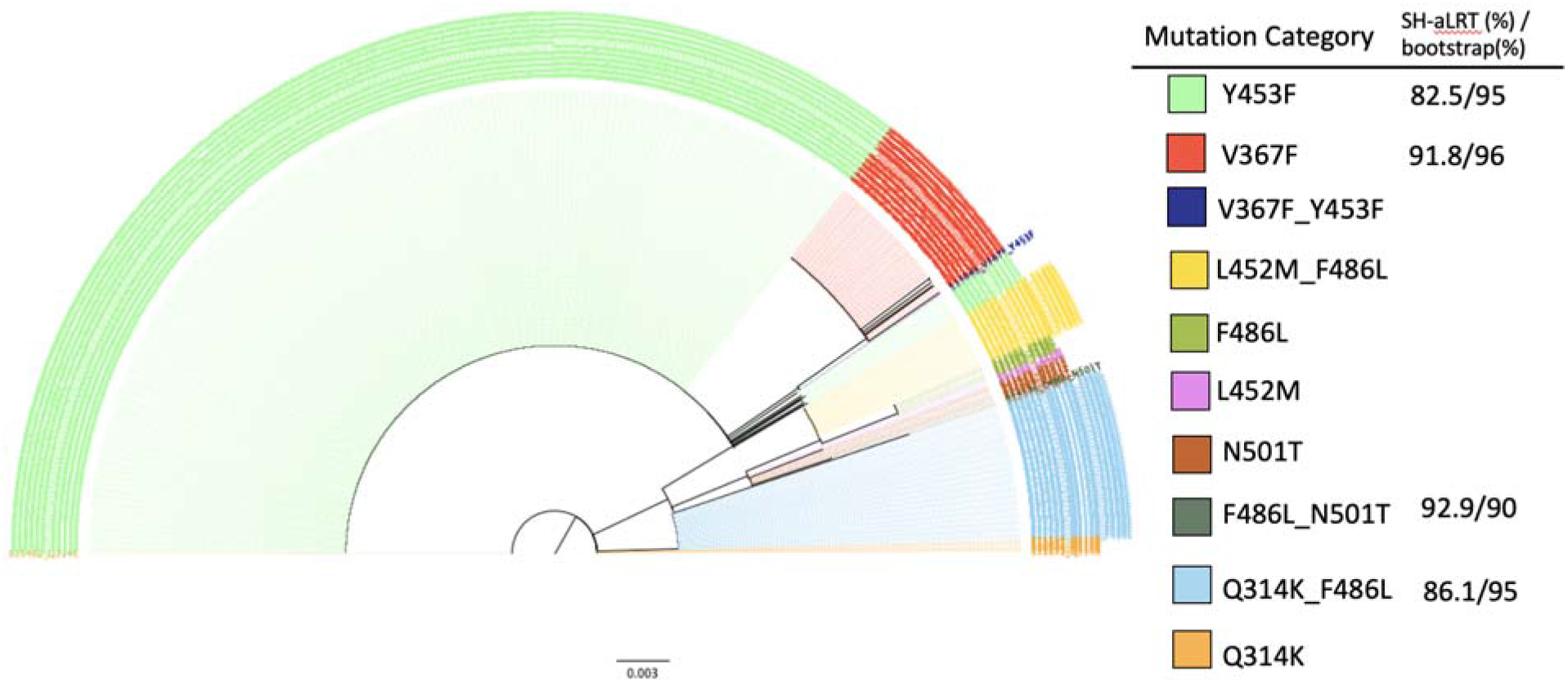
The phylogeny uses maximum likelihood with ultrafast bootstrapping till convergence and likelihood testing to generate the tree; shown if meeting high certainty threshold. Distinct, significant groupings were detected between all combinations of mutations, as opposed to a mixture of mutation type. Branch lengths to help determine differentiation or relationships are shown to scale at 0.003.

Bootstrap values for the plot completed early at 709 iterations due to a correlation coefficient at 0.996 as well as the absence of a new bootstrapped tree with better log-likelihood cutoffs. SH-like approximate likelihood ratio test^32^ testing was performed on each bootstrap. The bootstrap and Shimodaira–Hasegawa-like test values meeting the IQ-trees recommended threshold are shown in figure 4. Four of the 10 categories had branches meeting the strict threshold of 80% SH-like testing and 95% bootstrapping. The ultrafast bootstrap resampling and an additional SH-like branch test utilize imputed data to detect a significant difference from the original computed tree. Multiple full generations of the tree resulted in similar stratification of mutation categories. Bootstrapping takes about a 66% subset of a sequence’s positions to decrease time complexity, so the model will not contain certain mutations in some iterations. Since groupings generally have a one or two mutation difference max, there are slight differences from run to run in placement, but the observation of groupings remains true.

## 4. Discussion

The mutations outlined for SARS-CoV-2 show a pattern indicative of zoonotic transfer from humans to minks and back to humans.

The presence of F486L paired with either L452M or Q314K in humans and in minks indicates two separate transfer events between the species. These mutation pairs were observed in the Netherlands, but not elsewhere (table 1). The mutation F486L did not make a correlated jump from minks to humans until L452M, Q314K, or N501T was simultaneously present as a second mutation (figure 3). This illustrates the possibility that multiple mutations are required to preserve fitness and facilitate inter-species transfer, particularly in relation to the hosts ACE2 protein. Mutations in viruses have been described previously to occur in pairings for functional purposes^33^ and furthermore have been shown to have evolutionary relationships involving pairs of variants.

In Denmark, Y453F showed a pattern potentially arising from transmission to humans, then to minks, and back to humans again. Thousands of miles away in the US, isolated incidences of F486L, N501T, and V367F present evidence for the same type of transfer event, in the same species of mink. These data support the interpretation that paired mutations facilitate, or are required, for zoonotic transfer.

Although some submitted human sequences from the Netherlands do not have recorded collection dates, the submission timeline supports that the spread and transfer from minks to humans is occurring, as suggested by the submitters of the sequences,^35^ within the GISAID database^6,7^.

Current antibody-based therapeutics are focused on antibody binding to the SARS-CoV-2 spike protein RBD^36^. Similarly, spike protein antibody response-based vaccines may be dependent on the stability of the primary amino acid sequence in the RBD to maintain their ability to generate neutralizing antibody responses ^37^. It is therefore critical to understand the extent to which SARS-CoV-2 mutations are occurring in regions targeted by antibodies.

The extent to which mutations in the RBD could have a beneficial or deleterious effect on viral fitness, on RBD binding affinity to ACE2, and/or on infectivity is also not known. The mutation N501T does not appear to be spreading rapidly and may be showing decreased fitness in humans. Y453F however, now present in 629 human samples from Denmark beginning in June 2020, may be conferring increased viral fitness, potentially facilitating its spread into human populations. Our calculations also suggest that the N501T variant decreases protein stability and affinity for hACE2, but that the change is minimal. The pairing of certain mutations should be tested *in vitro* in the future. A single mutation could have a combinatory effect when paired with another, producing a completely different effect. The data suggest that two mutations together may be required for bi-directional zoonotic transfer to occur. While the number of cases with these pairings is not growing exponentially around the world, a third mutation could occur, further changing binding affinity and stability characteristics.

In Denmark, 629 human samples and 85 mink samples have shown the presence of Y453F. As this mutation may escape antibody neutralization,^38^ the emergence of this variant may increase resistance to monoclonal antibody therapy or convalescent sera therapy.

The evidence for SARS-CoV-2 zoonotic transfer using mutation analysis of human and other associate species that can carry the virus has helped identify how variants can arise. As there are multiple species that can be infected by SARS-CoV-2 to assist in this mutation potential,^39^ vigilant continual sequencing of the virus in the coming months and years is of high importance. Several species known to be infected by this virus have not been sequenced to the same extent, and although a large number of viruses isolated from minks were sequenced in the Netherlands and Denmark, as of this writing mink sequences from the United States are not publicly available. Tracking viral mutations isolated from minks and other farm animals that might later appear in humans is important. In Oregon for example, the location of the mink farm with SARS-CoV-2 outbreak has not been disclosed ^40^. Similarly, an outbreak in mink farms in Canada ^41^ has provided only 4 samples for analysis, all collected on December 4^th^, 2020. The mutation F486L has been found in only one of the four mink sequences available suggesting that these and other farms need to be carefully watched for the appearance of the second mutation. This would likely be either L452M or Q314K, now thought to enable the jump of the SARS-CoV-2 back to humans, as described in figure 3. Furthermore, although L452R has become increasingly common in humans with lineages B.1.427/429, this mutation differs from the position 452 mutation described in minks, L452M, indicating that it may have a role in viral transmission or infectivity.

Humans and minks have as much as 92% amino acid sequence similarity between their respective ACE2 receptor proteins (supplementary figure 1 & table 5). Of the residues of ACE2 that interact with the receptor binding domain of spike, mink and human are 83% similar while human and mice ACE2 are only 70% similar (supplementary table 5). Many different species have homologous ACE2, with a highly conserved binding site for the spike protein, demonstrating a wide range of potential hosts for a viral reservoir where new mutations may acquire new infectivity potential for humans ^42,43^. Based upon changes in affinity calculations when the human residues are substituted for mouse ACE2 residues, the affinity of ACE2 for spike decreases in most cases (supplemental table 5). However, when mink residues are substituted for human ACE2 residues, the affinity is minimally affected (except for G354H; supplemental table 5). This may explain why mice do not get infected by SARS-CoV-2 whereas the virus thrives in human and mink hosts.

Indeed, a novel artificial intelligence algorithm has shown that minks, along with bats, could be a reservoir of SARS-CoV-2 ^44^ and samples from cats, dogs, ferrets, hamsters, primates, and tree shrews demonstrate that all of these species have been infected with SARS-CoV-2 ^45^. Multiple factors contribute to zoonotic transfer including ACE2 expression level and close contact with the same or other species and therefore viral sequences of animals that may be in contact with humans is of great importance.

The consequence of observed mutations can be analyzed *in silico*, at least initially. Stability (expressed in terms of energy as kcal/mol), impact on RBD ACE2 binding or RBD and an antibody can be predicted. Additionally, prediction algorithms can determine the extent to which a single mutation or a combination of mutations could positively or negatively alter the function of a protein. These and other computational methods have already been used to assist in rapid vaccine design ^46^. Together, these methods can be utilized to identify optimal targets for vaccines and antibody therapeutics, ^47,48^ especially as selection pressure from the widespread use of these interventions increases the potential for virus escape ^13,49^.

## 5. Conclusion

Mutations in the RBD of SARS-CoV-2 have likely enabled zoonotic transfer beginning with the pandemic in China. We are now seeing zoonotic transfer most recently for minks in the Netherlands, Denmark, United States, and Canada. Whether these mutations become widespread and affect the efficacy of vaccines and monoclonal antibody cocktail therapeutics remains to be seen, but the potential exists for future mutations arising due to close contact between humans and a wide range of species. There is significant cause for concern that SARS-CoV-2 may evolve further with zoonotic transfer, which can involve various species, as it did at the origin of the outbreak. The emerging new variants acquired through such zoonotic transfer may facilitate human viral escape and the reduction in the efficacy of antibody-based vaccines and therapies. To help stop the pandemic, worldwide sequencing surveillance of samples from many species and correlating these with samples from humans will need to be performed as an early warning system for potential emergence of more virulent variants.

## Author Contributions

**Scott Burkholz:** Conceptualization, Methodology, Software, Visualization, Formal analysis, Writing - Original Draft, Writing - Review & Editing. **Suman Pokhrel, Benjamin R. Kraemer:** Methodology, Software, Visualization, Formal analysis, Writing - Original Draft, Writing - Review & Editing. **Richard T. Carback:** Software, Data Curation, Visualization, Formal analysis, Writing - Original Draft, Writing - Review & Editing **Daria Mochly-Rosen, Tom Hodge, Paul Harris, Serban Ciotlos, Lu Wang, CV Herst:** Formal analysis, Writing - Original Draft, Writing - Review & Editing. **Reid Rubsamen:** Formal analysis, Writing - Original Draft, Writing - Review & Editing, Supervision

## Data Availability

Sequence data not available to be distributed publicly per user agreement with GISAID.

The code used to discover mutations between humans and minks is available on gitlab. https://gitlab.com/flowpharma/papers/minkmutationanalysis2020

## Acknowledgements

The authors would like to thank scientists throughout the world that provided the SARS-CoV-2 sequences rapidly for on-going analyses during this pandemic.

---

Supplementary Table 1:

GISAID sequence citations with ID, authors, and originating laboratory are attached.

**Supplementary Table 2:**

| F486L |  |  | L452M |  |  | Y453F |  |  | N501T |  |  | V367F |  |  | Q314K |  |  |
| --- | --- | --- | --- | --- | --- | --- | --- | --- | --- | --- | --- | --- | --- | --- | --- | --- | --- |
| mutation | dStability | SNP | mutation | dStability | SNP | mutation | dStability | SNP | mutation | dStability | SNP | mutation | dStability | SNP | mutation | dStability | SNP |
| 1:F486F | 0.000 | 1 | 1:L452L | 0.000 | 1 | 1:Y453Y | 0.000 | 1 | 1:N501N | 0.000 | 1 | 1:V367V | 0.000 | 1 | 1:Q314Q | 0.000 | 1 |
| **1:F486L** | **0.716** | **1** | 1:L452F | 0.875 | 1 | **1:Y453F** | **0.695** | **1** | **1:N501T** | **0.915** | **1** | 1:V367I | 0.331 | 1 | 1:Q314L | 0.081 | 1 |
| 1:F486I | 0.746 | 1 | 1:L452R | 1.014 | 1 | 1:Y453H | 1.902 | 1 | 1:N501D | 1.081 | 1 | **1:V367F** | **0.336** | **1** | 1:Q314R | 0.649 | 1 |
| 1:F486Y | 0.996 | 1 | **1:L452M** | **1.116** | **1** | 1:Y453D | 2.484 | 1 | 1:N501S | 1.124 | 1 | 1:V367M | 0.449 | 1 | 1:Q314H | 1.049 | 1 |
| 1:F486V | 1.208 | 1 | 1:L452Q | 1.329 | 1 | 1:Y453N | 2.500 | 1 | 1:N501H | 1.687 | 1 | 1:V367L | 0.785 | 1 | 1:Q314E | 1.199 | 1 |
| 1:F486C | 1.941 | 1 | 1:L452W | 1.337 | 1 | 1:Y453C | 2.523 | 1 | 1:N501I | 3.430 | 1 | 1:V367A | 1.214 | 1 | **1:Q314K** | **1.262** | **1** |
| 1:F486S | 2.152 | 1 | 1:L452S | 1.462 | 1 | 1:Y453S | 2.794 | 1 | 1:N501K | 3.648 | 1 | 1:V367E | 1.278 | 1 | 1:Q314P | 1.640 | 1 |
| 1:F486M | 0.866 | 0 | 1:L452V | 1.465 | 1 | 1:Y453M | 1.479 | 0 | 1:N501Y | 33.473 | 1 | 1:V367G | 1.365 | 1 | 1:Q314I | 0.161 | 0 |
| 1:F486W | 1.442 | 0 | 1:L452H | 1.590 | 1 | 1:Y453W | 1.738 | 0 | 1:N501L | 0.482 | 0 | 1:V367D | 1.771 | 1 | 1:Q314F | 0.377 | 0 |
| 1:F486T | 1.628 | 0 | 1:L452I | 1.778 | 1 | 1:Y453Q | 2.118 | 0 | 1:N501C | 0.898 | 0 | 1:V367R | 0.615 | 0 | 1:Q314M | 0.401 | 0 |
| 1:F486Q | 1.655 | 0 | 1:L452P | 3.124 | 1 | 1:Y453T | 2.597 | 0 | 1:N501A | 0.967 | 0 | 1:V367Y | 0.663 | 0 | 1:Q314V | 0.522 | 0 |
| 1:F486E | 1.797 | 0 | 1:L452Y | 0.931 | 0 | 1:Y453A | 2.658 | 0 | 1:N501M | 1.493 | 0 | 1:V367W | 0.863 | 0 | 1:Q314Y | 0.546 | 0 |
| 1:F486R | 1.940 | 0 | 1:L452A | 1.245 | 0 | 1:Y453E | 2.964 | 0 | 1:N501G | 1.872 | 0 | 1:V367K | 0.945 | 0 | 1:Q314W | 0.775 | 0 |
| 1:F486H | 1.942 | 0 | 1:L452C | 1.326 | 0 | 1:Y453V | 3.565 | 0 | 1:N501Q | 2.025 | 0 | 1:V367Q | 1.053 | 0 | 1:Q314T | 0.777 | 0 |
| 1:F486A | 1.955 | 0 | 1:L452N | 1.408 | 0 | 1:Y453G | 3.577 | 0 | 1:N501E | 2.866 | 0 | 1:V367C | 1.169 | 0 | 1:Q314A | 0.934 | 0 |
| 1:F486N | 2.001 | 0 | 1:L452E | 1.475 | 0 | 1:Y453R | 5.515 | 0 | 1:N501V | 3.857 | 0 | 1:V367T | 1.288 | 0 | 1:Q314G | 1.008 | 0 |
| 1:F486K | 2.120 | 0 | 1:L452K | 1.517 | 0 | 1:Y453I | 7.582 | 0 | 1:N501R | 7.724 | 0 | 1:V367S | 1.399 | 0 | 1:Q314C | 1.015 | 0 |
| 1:F486D | 2.288 | 0 | 1:L452D | 1.668 | 0 | 1:Y453P | 10.805 | 0 | 1:N501F | 12.759 | 0 | 1:V367N | 1.507 | 0 | 1:Q314N | 1.041 | 0 |
| 1:F486G | 2.371 | 0 | 1:L452G | 1.847 | 0 | 1:Y453K | 11.958 | 0 | 1:N501P | 102.865 | 0 | 1:V367H | 1.705 | 0 | 1:Q314S | 1.169 | 0 |
| 1:F486P | 2.518 | 0 | 1:L452T | 2.451 | 0 | 1:Y453L | 13.639 | 0 | 1:N501W | 105.796 | 0 | 1:V367P | 3.039 | 0 | 1:Q314D | 1.212 | 0 |

**Supplementary Table 3:**

| F486L |  |  | Y453F |  |  | N501T |  |  | L452M |  |  |
| --- | --- | --- | --- | --- | --- | --- | --- | --- | --- | --- | --- |
| mutation | daffinity | SNP | mutation | daffinity | SNP | mutation | daffinity | SNP | mutation | dAffinity | SNP |
| 1:F486Y | -0.578 | 1 | **1:Y453F** | **-0.080** | **1** | 1:N501H | -0.250 | 1 | 1:L452H | -0.082 | 1 |
| 1:F486F | 0.000 | 1 | 1:Y453Y | 0.000 | 1 | 1:N501N | 0.000 | 1 | **1:L452M** | **-0.038** | **1** |
| **1:F486L** | **2.796** | **1** | 1:Y453N | 0.559 | 1 | 1:N501D | 0.278 | 1 | 1:L452P | -0.010 | 1 |
| 1:F486I | 2.912 | 1 | 1:Y453S | 0.588 | 1 | **1:N501T** | **0.872** | **1** | 1:L452F | -0.005 | 1 |
| 1:F486V | 4.245 | 1 | 1:Y453H | 0.598 | 1 | 1:N501S | 2.281 | 1 | 1:L452Q | -0.004 | 1 |
| 1:F486C | 4.583 | 1 | 1:Y453C | 0.606 | 1 | 1:N501Y | 2.286 | 1 | 1:L452S | -0.003 | 1 |
| 1:F486S | 5.042 | 1 | 1:Y453D | 0.687 | 1 | 1:N501I | 13.551 | 1 | 1:L452L | 0.000 | 1 |
| 1:F486W | -0.296 | 0 | 1:Y453W | -0.429 | 0 | 1:N501K | 15.768 | 1 | 1:L452I | 0.006 | 1 |
| 1:F486M | 0.776 | 0 | 1:Y453K | -0.411 | 0 | 1:N501Q | -1.315 | 0 | 1:L452V | 0.014 | 1 |
| 1:F486Q | 0.856 | 0 | 1:Y453M | 0.293 | 0 | 1:N501E | -0.826 | 0 | 1:L452W | 0.040 | 1 |
| 1:F486H | 1.433 | 0 | 1:Y453E | 0.375 | 0 | 1:N501M | 0.797 | 0 | 1:L452R | 0.195 | 1 |
| 1:F486E | 1.989 | 0 | 1:Y453Q | 0.388 | 0 | 1:N501V | 1.205 | 0 | 1:L452E | -0.165 | 0 |
| 1:F486R | 2.077 | 0 | 1:Y453I | 0.496 | 0 | 1:N501C | 1.229 | 0 | 1:L452D | -0.013 | 0 |
| 1:F486K | 3.285 | 0 | 1:Y453L | 0.512 | 0 | 1:N501L | 2.066 | 0 | 1:L452G | -0.010 | 0 |
| 1:F486N | 3.786 | 0 | 1:Y453P | 0.532 | 0 | 1:N501P | 2.123 | 0 | 1:L452A | -0.004 | 0 |
| 1:F486T | 4.312 | 0 | 1:Y453T | 0.538 | 0 | 1:N501A | 2.721 | 0 | 1:L452N | -0.003 | 0 |
| 1:F486D | 4.431 | 0 | 1:Y453V | 0.561 | 0 | 1:N501G | 3.323 | 0 | 1:L452T | -0.001 | 0 |
| 1:F486P | 5.072 | 0 | 1:Y453G | 0.593 | 0 | 1:N501F | 49.293 | 0 | 1:L452Y | 0.016 | 0 |
| 1:F486A | 5.334 | 0 | 1:Y453A | 0.600 | 0 | 1:N501R | 64.572 | 0 | 1:L452C | 0.027 | 0 |
| 1:F486G | 5.827 | 0 | 1:Y453R | 25.426 | 0 | 1:N501W | 306.442 | 0 | 1:L452K | 0.062 | 0 |

**Supplementary Table 4:**

| Variant | Within 4.5Å<br>of ACE2? | PROVEAN | SIFT |
| --- | --- | --- | --- |
| F486L | Yes | Neutral | Tolerated |
| L452M | No | Neutral | Tolerated |
| Q314K | No | Neutral | Tolerated |
| Y453F | Yes | Neutral | Tolerated |
| N501T | Yes | Neutral | Tolerated |
| V367F | No | Neutral | Tolerated |

Supplementary Table 5: File is attached.

**Supplementary Figure 1:**
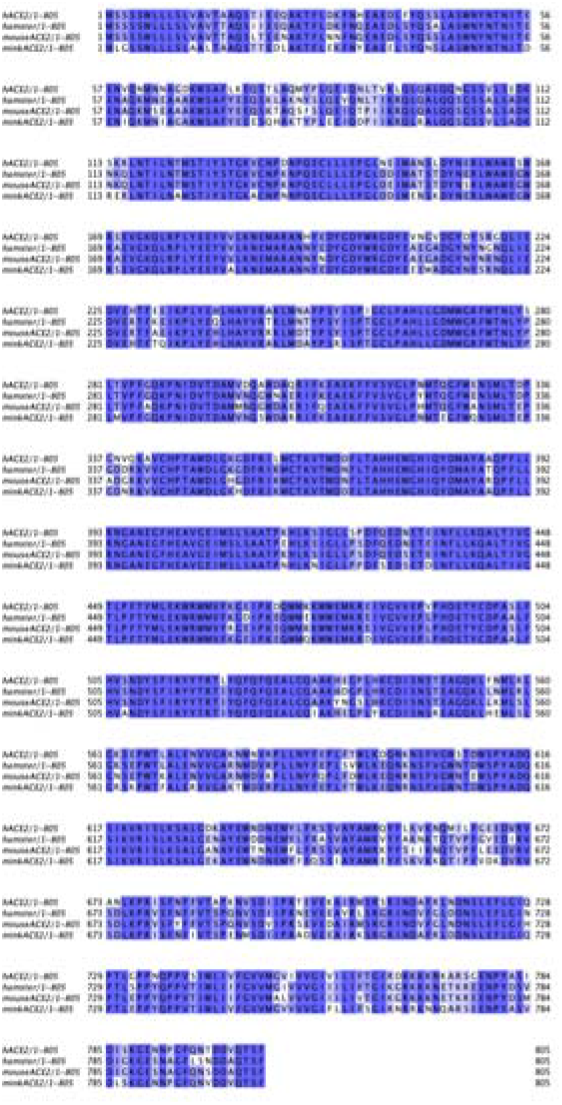

